# KLF5-regulated extracellular matrix remodeling secures biliary epithelial tissue integrity against cholestatic liver injury

**DOI:** 10.1101/2022.01.25.477619

**Authors:** Minami Yamada, Hajime Okada, Masatsugu Ema, Yamato Kikkawa, Atsushi Miyajima, Tohru Itoh

## Abstract

Tubular epithelial tissues in the body play fundamental roles as infrastructure constituting conduits to transport various types of biological fluids, for which contiguous and integrated epithelial tissue structures should be maintained continuously and even under stressed conditions. Compared to tissue morphological processes that take place during ontogeny, the mechanisms whereby tubular epithelial tissues maintain their structural integrity in adulthood remains largely unclear. Here, we show that the transcription factor Klf5 is crucial for maintaining the biliary epithelial integrity in tissue remodeling processes induced under cholestatic injury conditions in the adult liver. Loss of Klf5 in the biliary epithelia led to tissue collapse *in vivo* in injured mouse livers, as well as *in vitro* in bile ductular organoids in a tissue-autonomous manner and independent of cell proliferation. Klf5 regulated cell junction organization and cell adhesion, along with extracellular matrix remodeling around the expanding biliary epithelia through deposition of Lamb3-containing laminin complexes. Targeting the Lamb3 expression in biliary epithelia in mice recapitulated the tissue collapse phenotype. Together, our results highlight a novel mechanism whereby the epithelial tissue maintains its integrity while undergoing unstable structural transformation.

## Main

The bile duct in the liver is a typical tubular epithelium composed of biliary epithelial cells (BECs; also known as cholangiocytes), forming a tree-like structure throughout the organ to constitute the drainage system for the hepatocyte-borne bile, a highly cytotoxic fluid whose leakage causes detrimental effects on the liver tissue and the organism. Under a variety of liver injury conditions in human diseases and in animal models, the biliary epithelia undergo dynamic tissue expansion and structural remodeling^1^, also referred to as the ductular reaction^2,3^, to form auxiliary conduit networks to ensure proper bile excretion^4,5^. The biliary epithelial tissue remodeling/ductular reaction is an adaptive response involving arborization and expansion of the branches so that they extend toward the parenchymal injury region^1,6,7^, and plays multifaceted roles in liver regeneration and pathophysiology^8,9^.

In the course of studies elucidating the molecular mechanisms of the ductular reaction, we recently identified Krüppel-like factor 5 (Klf5) as a transcription factor that is prominently expressed in BECs and critical for their regulation^10^. Thus, the liver epithelial-specific Klf5 knockout mice (Klf5-LKO mice) exhibited significant defects in the ductular reaction in models of a subtype of liver injury characterized by severe cholestasis, leading to an increased mortality rate. While the defects in the ductular reaction well correlated with suppression of BEC proliferation, remarkably, we also noticed the presence of isolated cells and cell clusters that were positive for the biliary marker but were clearly separated from the original contiguous biliary tissue structure in injured LKO livers (Figure 1A). This implies a potential additional role of Klf5 in the maintenance of tissue integrity other than cell proliferation.

**Figure 1.**
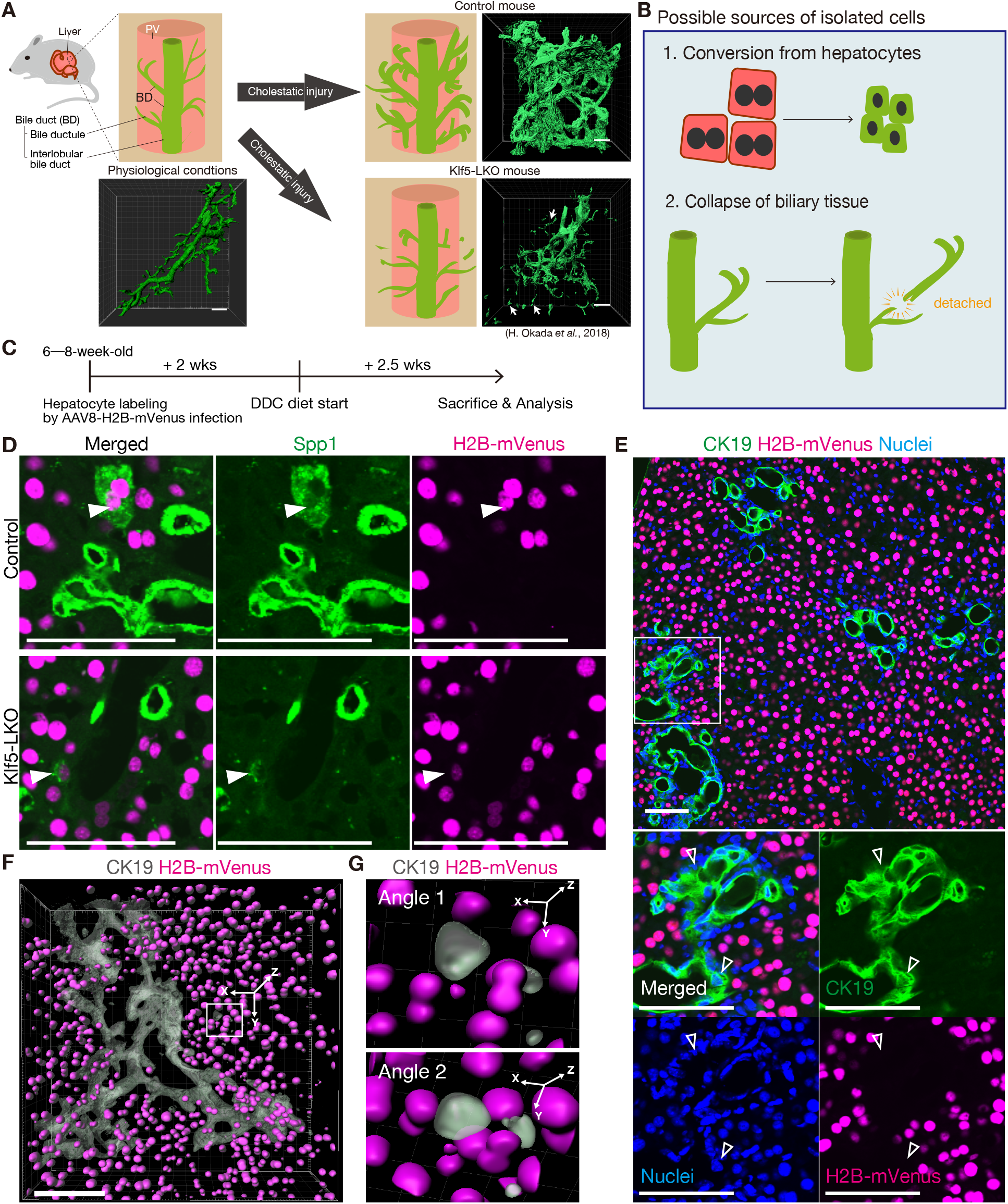
Aberrantly isolated BECs in the cholestatic Klf5-LKO mouse livers are not of hepatocyte origin. (A) Schematic illustration and representative images for bile duct (BD; shown in green) morphologies before and after cholestatic liver injury. In the schematic illustration, portal veins (PVs) are indicated in pink. The photographs are reproduced from Okada, H., et al. (ref. 10) and Yamada, M., et al. (ref. 24). The arrows point to the isolated CK19^+^ biliary epithelial cells and clusters observed specifically in the cholestatic liver of Klf5-LKO mice. Scale bars, 100μm. (B) Working hypothesis on the origin of isolated biliary epithelial cells and clusters. (C) Experimental scheme of hepatocytes lineage-tracing experiments via rAAV2/8-H2B-mVenus infection. (D) Representative 2D images of the rAAV2/8-H2B-mVenus-infected liver sections after DDC administration. H2B-mVenus signals (magenta) are shown with immunostaining signals for Spp1 (green). The white arrowheads point to Spp1^+^ H2B-mVenus^+^ “biliary-like hepatocytes” that are characteristically observed in cholestatic livers and are known to originate from hepatocytes. Scale bars, 100 μm. (E) Representative 2D images of the rAAV2/8-H2B-mVenus-infected liver sections from the control mice after DDC administration. H2B-mVenus signals (magenta) are shown with immunostaining signals for CK19 (green) and nucleus counterstaining with Hoechst (blue). A region indicated by a white box in the upper panel is magnified in the lower panels. The blank arrowheads indicate CK19^+^ BECs. Scale bars, 100 μm. (F and G) Representative 3D images of the rAAV2/8-H2B-mVenus-infected Klf5-LKO liver sections after DDC administration, showing biliary epithelial tissue structure (CK19 immunostaining, gray) and H2B-mVenus signals (magenta). The 3D images were processed as explained in Supplementary Figure 1 so that H2B-mVenus signals out of the Z-plane where the isolated CK19^+^ cells reside are eliminated. In G, magnified images corresponding to the region indicated by the white rectangle in F are shown. The upper panel (Angle 1) is from almost the same angle as in F, and the lower panel (Angle 2) is a rotated view, confirming that the cells are completely separated from the biliary tree. Scale bar, 100 μm.

It has become evident in recent years that the liver epithelial tissue, consisting of hepatocytes and BECs, exhibits substantial levels of lineage plasticity, in that the two types of cells can undergo bidirectional conversion with each other in response to various types of injuries^9^. In particular, lineage conversion of hepatocytes to biliary-type cells (also known as ductal metaplasia) is typically induced upon cholestatic liver injury or in animal models of ductopenia^11–13^. Given that the Klf5-LKO mice exhibit massively exacerbated cholestatic injury phenotypes compared to control mice under corresponding disease conditions^10^, we first speculated that the aforementioned biliary marker-positive cell isolates observed in these mice might be derived from aberrant transformation of hepatocytes (Figure 1B). As the loss of the Klf5 gene is achieved not only in BECs but also in hepatocytes (owing to Alfp-Cre–mediated deletion in fetal liver progenitor cells for both lineages), hepatocyte-derived biliary-type cells should have somehow been defective in forming an authentic and contiguous biliary epithelial tissue structure.

To test this hypothesis, we set out to perform an *in vivo* lineage tracing analysis of hepatocytes in the background of the Klf5-LKO mice. As conventional cell labeling approaches based on the Cre/loxP-mediated marker gene expression were not compatible, we made use of a histone H2B-fused fluorescent reporter whose expression in the cell nuclei can be stably maintained over a certain period of time and upon cell differentiation^14^. We adopted the hepatotropic adeno-associated viral vector system with type-VIII capsid (AAV8)^15^ to express H2B-fused mVenus (H2B-mVenus) proteins exclusively in hepatocytes but not in BECs or any other cell types in the mouse liver (Figure 1C). When mice were subjected to this hepatocyte labeling system followed by a cholestatic liver injury regimen using a 3,5-diethoxycarbonyl-1,4-dihydrocollidine (DDC)-containing diet, H2B-mVenus^+^ hepatocytes that had become positive for the biliary marker Spp1/osteopontin were detected around the PV in both the Klf5-LKO and the control livers (Figure 1D). Nevertheless, H2B-mVenus signals did not coincide with the authentic biliary marker CK19, which is fully consistent with the mode of acquisition of “biliary-like” phenotypes in hepatocytes upon DDC-induced cholestasis as previously reported^11,15^ (Figure 1E). Importantly, the isolated CK19^+^ cells in the Klf5-LKO mouse livers were negative for the H2B-mVenus expression (Figure 1F and G, and Supplementary Figure S1). These results indicate that the isolated BEC clusters in the cholestatic Klf5-LKO mice did not originate from hepatocytes, rather likely representing degradants of the pre-existing biliary epithelial tissue (Figure 1B).

To characterize whether and how the collapse of the biliary epithelial tissue occurred, we sought to develop an *in vitro* model system with which time-lapse monitoring of the process could be achieved. Conventional biliary organoids produced by Matrigel embedding exhibit cyst-like morphology^16–18^ and likely represent the interlobular bile duct (see Figure 1A). In contrast, the culture of BECs in collagen gels can reproduce the growth of the biliary epithelial tissue with branched 3D morphology^19^, which more faithfully represents remodeling bile ductules in the ductular reaction. As BECs isolated from the adult Klf5-LKO mice, which had already lost the Klf5 expression, were not proliferative *per se* and could not be maintained under culture conditions (data not shown), we constructed a BEC line where inducible knockout of Klf5 could be achieved. Thus, BECs were first isolated from *Klf5 flox/flox* mice and cultured, and a tamoxifen-inducible variant of the Cre recombinase, CreERT2, was introduced together with the mVenus fluorescent reporter by retroviral gene transfer (Supplementary Figure S2A, and see Methods). These cells (*Klf5 flox/flox;LTR-CreERT2-IRES-mVenus*, hereafter referred to as Klf5-iKO cells) were able to continuously grow and extend branches in collagen gels *in vitro*, reminiscent of the reacting ductules observed *in vivo* (Figure 2A). When conditional deletion of *Klf5* was subsequently induced in those *in vitro*-formed ductular structures by 4-hydroxytamoxifen (4-OHT), the branched tissue structure began to collapse, resulting in the appearance of separated cells and clusters (Figure 2, B−D, and Supplementary Video). We confirmed that *Klf5* expression was almost completely depleted at 24 h after 4-OHT addition prior to the induction of tissue collapse (Supplementary Figure S2B) and that 4-OHT itself did not cause any effect (Supplementary Figure S2C), supporting the notion that the collapse occurred due to the loss of Klf5. The collapse was induced both at the tip and in the middle of the biliary branches, and the detached cells displayed a round shape, as was observed *in vivo* in the knockout mouse liver (Figure 1, F and G). Notably, the collapse was induced in a biliary epithelium-autonomous manner in the absence of any other type of liver cells. Taken together, our success in simulating the ductular structure and its collapse *in vitro* led to the identification of a novel tissue-intrinsic mechanism governed by Klf5 whereby the biliary tissue maintains its epithelial structural integrity under expanding and remodeling conditions.

**Figure 2.**
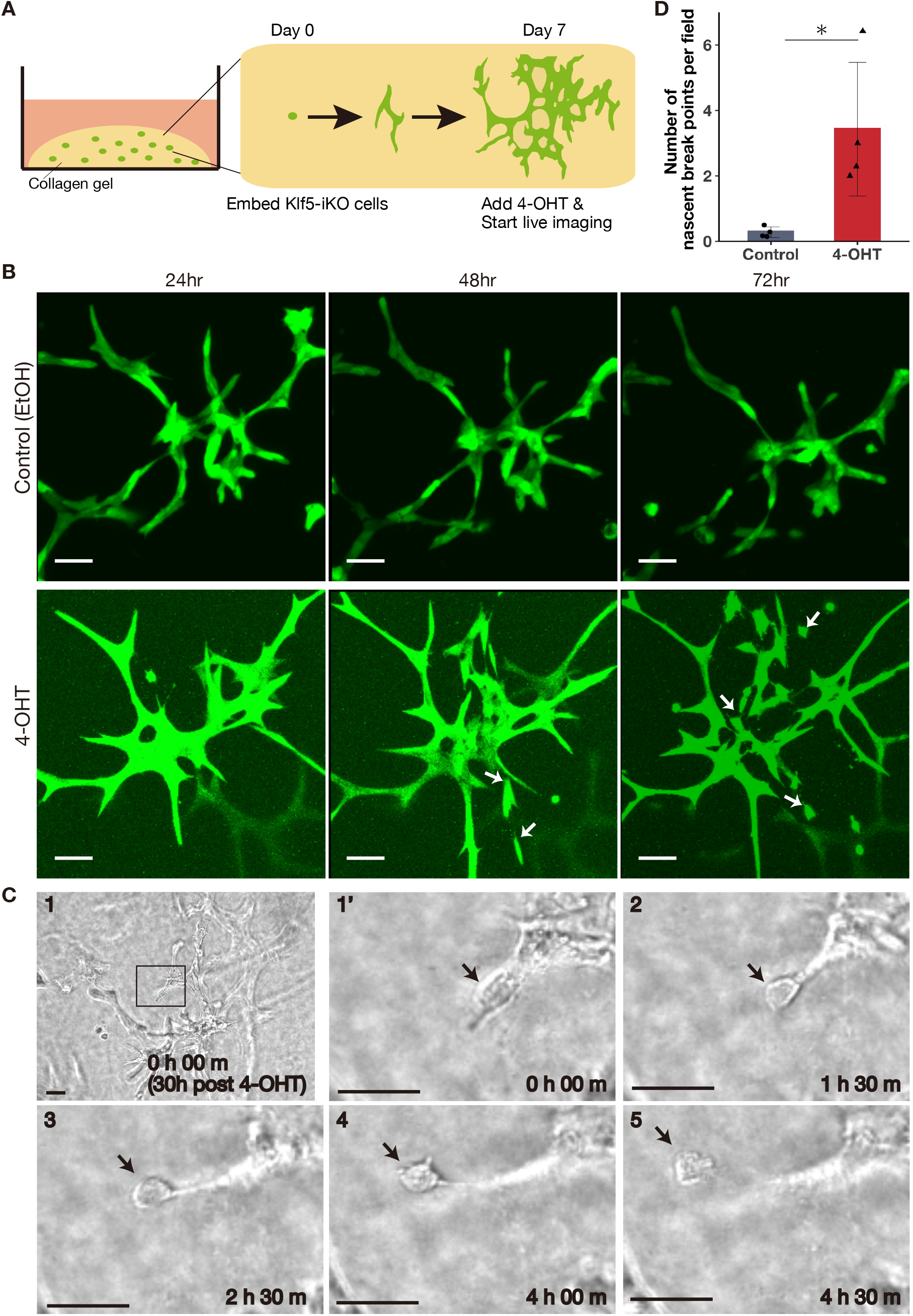
Loss of Klf5 leads to collapse of the expanding biliary epithelial tissue *in vitro*. (A) Schematic diagram of the *in vitro* culture system. Tamoxifen-inducible Klf5 knockout BEC line (Klf5-iKO cell line, see Supplementary Figure 2A; shown in green) was embedded in a collagen gel (yellow) and cultured for 7 days to grow branched biliary epithelial organoids. The organoids were then treated with or without 4-hydroxytamoxifen (4-OHT) and subjected to live imaging analyses. (B) Representative time-lapse mVenus fluorescence images demonstrating the biliary organoids at 24, 48, and 72 h after administration of 4-OHT (bottom panels) or EtOH as a control (top panels). The arrows indicate detached cell clusters. Scale bars, 100 μm. (C) Representative time-lapse bright-field images demonstrating the collapse of the biliary epithelial structure after 4-OHT administration. Panel 1 shows a lower magnified image for biliary organoids, and Panels 1’ and 2–6 show magnified images for the region indicated as a rectangle in panel 1 corresponding to a tip region of a biliary branch where a cell cluster was going to be detached. Elapsed time intervals after 4-OHT administration are indicated in hours (h) and minutes (m). Black arrows point to a cell cluster to be detached. Scale bars, 50 μm. (D) Quantification of the biliary epithelial tissue collapse. The numbers of nascent break points formed between a detached cell cluster and the biliary branching structure were counted in time-lapse images covering from 24 to 48 h after EtOH (control) and 4-OHT administration (*n* = 4 culture wells per condition), and data represent the mean ± S.D. Three to seven views per each well were observed. Dots in the graph indicate the nascent break point numbers in each well normalized to view numbers. The p-value was calculated using the two-sided Mann-Whitney U test. An asterisk indicates that the p-value is <0.05.

To unravel the mode of action of Klf5, RNA sequencing (RNA-seq) analysis was performed using the *in vitro* model system undergoing the course of tissue collapse upon 4-OHT treatment. The number of differentially expressed genes (DEGs) in the degenerating tissues upon *Klf5* deletion compared to the expanding control tissues increased with the lapse of time (24 h vs. 12 h; Supplementary Figure S3, A and B), and their associated gene ontology (GO) terms included “cell-substrate adhesion,” “extracellular structure organization,” “cell junction organization,” and “regulation of cell adhesion” (Figure 3A), implicating Klf5 in regulating the cell–extracellular matrix (ECM) interaction to maintain biliary epithelial tissue integrity.

**Figure 3.**
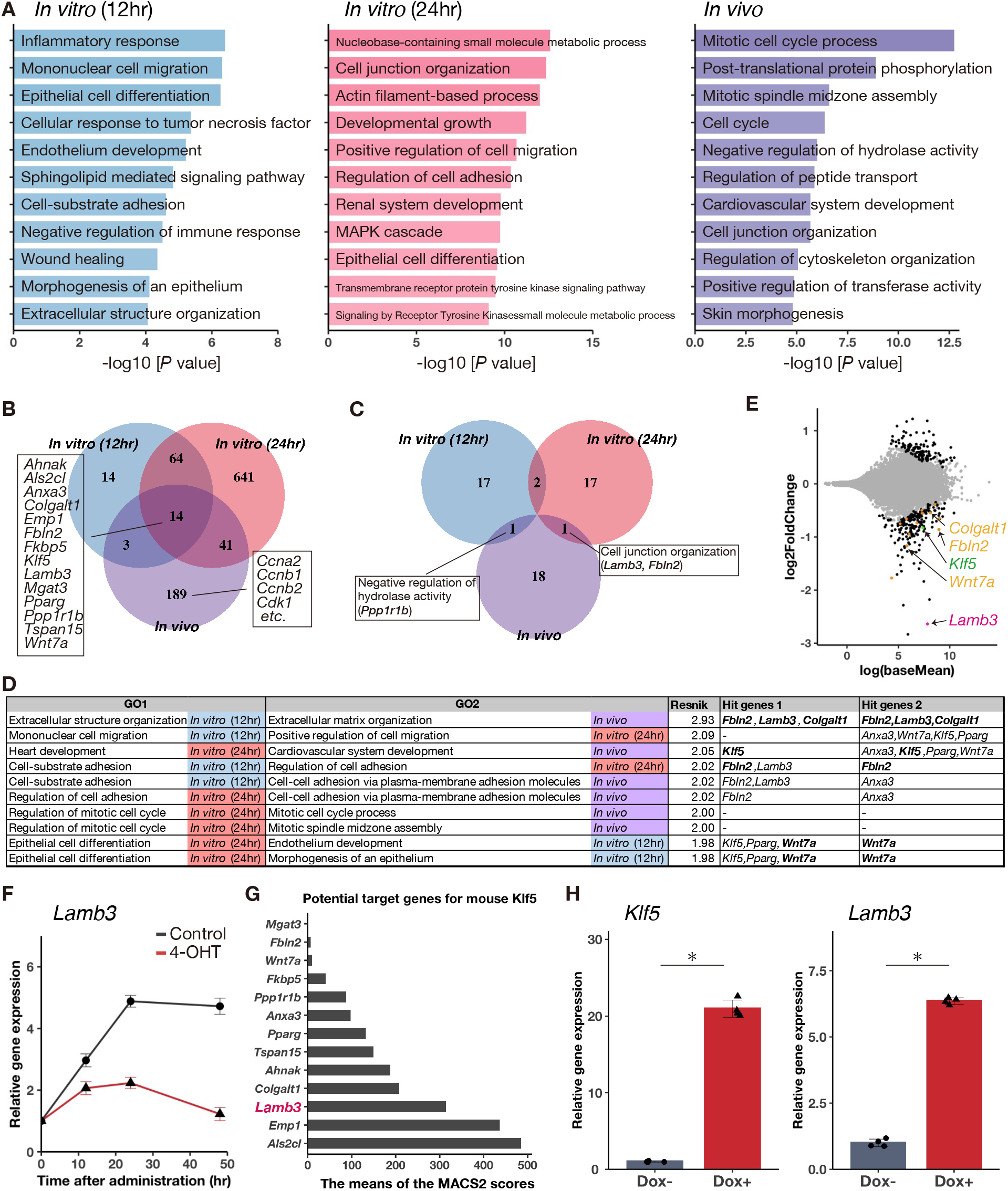
Exploring essential genes under the Klf5-mediated regulation for maintenance of a bile duct structure. (A) GO enrichment analysis of DEGs upon *Klf5* deletion in BECs using Metascape. RNA-seq data obtained from the *in vitro* culture system at 12 and 24 h after 4-OHT addition (vs. EtOH control) and those from the *in vivo* model in the Klf5-LKO vs. control mice under the DDC injury condition (ref. 10) were analyzed. Top 20 GO terms are shown. (B) Venn diagram showing the number and overlap of downregulated DEGs upon *Klf5* deletion among the three comparisons. Fourteen genes, including *Klf5*, were commonly downregulated in all the three comparisons, while expression of cell cycle-related genes was affected only in the *in vivo* condition. (C) Venn diagram showing the overlap of top 20 GO terms enriched by *Klf5* deletion among the three comparisons. (D) Extracting highly similar GO terms among the three comparisons in Figure 3A using NaviGO. Hit genes 1 and 2 indicate the “commonly downregulated 14 genes” shown in Figure 3B that were identified as responsible genes for GO1 and GO2, respectively, where those genes that are common between the two are shown in bold. (E) MA plot showing genes differentially expressed in BECs from the Klf5-LKO and control mouse livers under the DDC injury condition (ref. 10). Those common responsible genes identified and shown in bold in Figure 3D are highlighted in yellow color. The other DEGs and non-DEGs are shown in black and grey, respectively. (F) Time course of the *Lamb3* expression level in the *in vitro* culture system of the Klf5-iKO cells. The grey and red lines indicate control (EtOH)- and 4-OHT–treated samples, respectively. *n* = 4 biological replicates per time point for each of the treatment, and data represent the mean ± S.D. (G) The means of MACS2 scores (http://chip-atlas.org) calculated for potential target genes of mouse Klf5. (H) Expression levels of *Klf5* (left) and *Lamb3* (right) in the *in vitro* culture system of BECs with Klf5 overexpression. Tet-inducible Klf5-overexpressing BEC line was cultured for 7 d and then stimulated by Dox administration for 72 h. Data represent the mean ± S.D. of *n* = 4 biological replicates per condition. The p-value was calculated using the two-sided Mann-Whitney U test. An asterisk indicates that the p-value is <0.05.

To identify critical molecules and pathways functioning downstream of Klf5, we narrowed down the candidate target genes by combining the results of the present RNA-seq with those obtained previously using primary BEC samples isolated from the livers of Klf5-LKO and control mice under DDC-induced cholestatic injury^10^. We identified 14 significantly downregulated genes whose expression was consistently reduced in both *in vitro* and *in vivo* conditions, including several genes that are known to be related to cell junction organization and cell–ECM interactions (Figure 3B). Interestingly, while the loss of Klf5 *in vivo* in the knockout mice significantly affected cell cycle progression (Figure 3A) with reduced expression of relevant genes such as those encoding cyclins and *Cdk1* in BECs^10^, expression of these genes was not reduced in the ductular organoid culture *in vitro* (Figure 3B). The EdU incorporation assay confirmed that the loss of Klf5 did not affect the cell cycle progression *in vitro* (Supplementary Figure S3, C and D), suggesting that the biliary collapse was not a mere secondary event annexed to the cell cycle blockade but rather induced independently of cell proliferation.

Comparison of GO terms related to the three different sample conditions failed to extract any common biological processes (Figure 3C). Nevertheless, computing the similarity of strongly affected GO terms^20^ (top 20) among the three conditions led to the identification of highly similar terms together with common responsible genes, including *Fbln2, Lamb3, Colgalt1*, and *Wnt7a* (Figure 3D). Among these genes, we particularly focused on *Lamb3*, which encodes the β3 subunit of the ECM component laminin proteins, as it was among the most drastically reduced genes *in vivo* in the Klf5-deficient liver under cholestatic conditions (Figure 3E). Quantitative gene expression analysis demonstrated that the expression of *Lamb3* was also substantially suppressed *in vitro* upon the inducible loss of Klf5 (Figure 3F). *In silico* analysis of public ChIP-seq data^21^ showed that Klf5 is bound close to the transcriptional start sites of *Lamb3* and *LAMB3* in mouse and human genomes, respectively (Figure 3G, and Supplementary Figure S3E–G). Moreover, overexpression of Klf5 was capable of inducing *Lamb3* expression in BECs (Figure 3H), as has been reported in a chondrogenic cell line^22^. Together, these results implicate Lamb3 as a direct and functionally relevant target of Klf5 responsible for the biliary epithelial tissue maintenance.

Laminins are heterotrimeric ECM proteins composed of α, β and γ subunits, and serve as major components of the basal lamina. The *Lamb3*-encoded laminin β3 subunit, together with the α3 and γ2 subunits (encoded by Lama3 and Lamc2, respectively), forms laminin-332, which plays a major role in skin epithelium adherence^23^. In the liver, laminin-332 is distributed around the outer layer of the bile ducts, with its function being unidentified^24,25^. Intriguingly, our previous study indicated that DDC administration in mice led to a significant increase in mRNA expression levels of *Lama3* and *Lamb3* in BECs^10^, which prompted us to further investigate the spatial expression profiles of the laminin-332 components by immunostaining of liver tissue sections.

Intrahepatic bile ducts can be roughly divided into two morphologically distinct epithelial tissue structures, namely, interlobular bile ducts and bile ductules^15^ (Figure 4A); under physiological conditions in the absence of liver injury, the deposition of laminin-332 components is restricted only around the former but not the latter biliary epithelia^24^. Upon administration of DDC, expression of Lamb3 proteins became substantially expanded so that they accumulated around the entire biliary epithelial tissue, including the ductules that extended and dilated around the PV (Figure 4B), in line with upregulation at the mRNA level^10^. However, the deposition of Lamb3 was completely suppressed in the Klf5-deficient liver tissue, confirming a genetic relationship between Lamb3 and Klf5 (Figure 4C). Concomitant suppression of deposition of the Lama3 subunit was also observed around the bile ducts (Supplementary Figure S4A). As another cholestatic liver injury model, we also employed knockout mice for the ATP-binding cassette subfamily B member 4 (*Abcb4*) gene^26^, representing a hereditary cholestatic disorder known as progressive familial intrahepatic cholestasis type 3 in humans. The ductular reaction occurs in the liver of Abcb4 KO mice, which can be suppressed by the loss of Klf5 in the biliary epithelial tissue in the Abcb4 KO;Klf5-LKO double-knockout mice^10^. Immunostaining analyses revealed that Lama3 and Lamb3 were accumulated around the entire biliary epithelia, including ductules, in the Abcb4 KO liver; this accumulation was also suppressed in the liver of Abcb4 KO;Klf5-LKO mice concomitantly with diminished ductular reaction (Figure 4D, and Supplementary Figure S4B).

**Figure 4.**
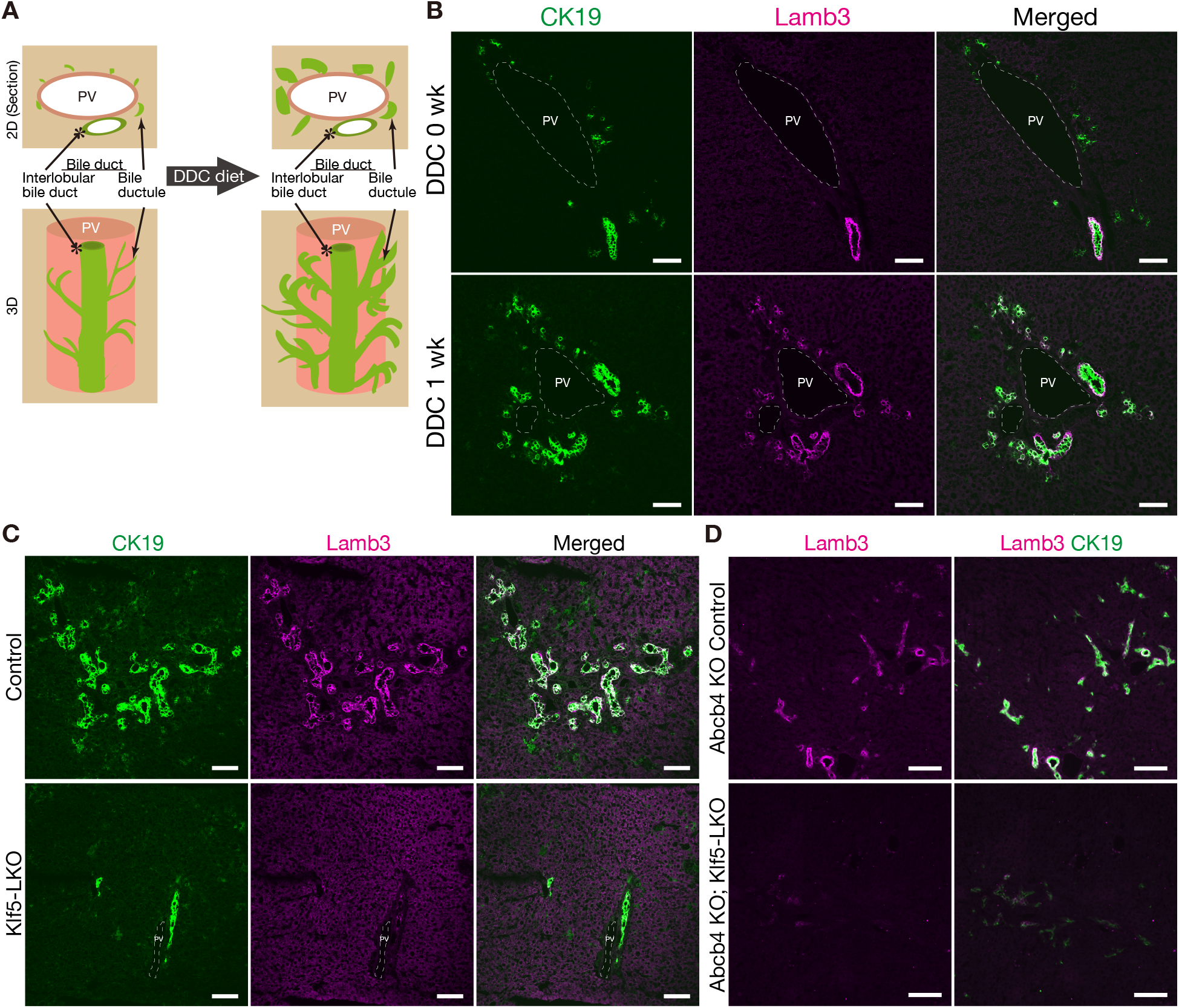
Deposition of Lamb3 is induced around the expanding biliary epithelia in cholestatic mouse livers. (A) Schematic illustration of the bile duct morphology and substructure under the normal (left) and DDC diet-induced cholestatic injury conditions (right). PV, portal vein. (B) Representative images of immunostaining for CK19 (green) and Lamb3 (magenta) in liver sections of wild-type mice before (0 wk) and 1 week after DDC treatment. Scale bars, 100 μm. (C) Representative images of immunostaining for CK19 (green) and Lamb3 (magenta) in liver sections of control (*Klf5 fl/fl*) and Klf5-LKO mice after 3 weeks of DDC administration, indicating the loss of Lamb3 expression in the Klf5-LKO mouse liver at the protein level. Scale bars, 100 μm. (D) Representative images of immunostaining for Lamb3 (magenta) and CK19 (green) in liver sections of *Abcb4* KO control (Abcb4 KO; *Klf5 fl/fl)* and *Abcb4* and *Klf5* double-knockout (Abcb4-KO; Klf5-LKO) mice at 8 weeks after birth. Scale bars, 100 μm.

To clarify the role of Lamb3 in the biliary remodeling, we used liver epithelial-specific *Lamb3* conditional knockout mice (Lamb3-LKO mice). The Lamb3-LKO mice followed the Mendelian ratio at birth, developed normally, and exhibited no defects in the hepatobiliary system under physiological conditions despite complete absence of Lamb3 expression in the liver^24^. Consistent with the assumption that Lamb3 is a downstream target of Klf5, expression of Klf5 in the biliary epithelial tissue was not affected in the Lamb3-LKO liver (Supplementary Figure S5A).

We then applied the DDC-induced cholestatic injury to the mutant mice, where Lamb3 proteins were still completely absent in the liver (Supplementary Figure S5B) with no effect on Klf5 expression (Supplementary Figure S5C). Intriguingly, expression of Lama3 was severely suppressed in the Lamb3-deficient liver (Supplementary Figure S5D) and that of Lamc2 also appeared to be decreased (Supplementary Figure S5E). These results are consistent with the notion that the heterotypic complex formation precedes secretion and extracellular deposition of laminin complexes^27^ and indicate that the functional laminin-332 complex is lost in the liver of the Lamb3 knockout mice. In contrast with the Klf5-LKO mice, however, the biliary branches continued to expand in the Lamb3-LKO mice under DDC injury conditions (Figure 5A). Indeed, quantification of the CK19^+^ area in the liver sections revealed that the level of the ductular reaction was suppressed only marginally in the absence of Lamb3 (Figure 5B). Immunostaining analysis for Ki67 revealed that proliferation of BECs was not affected in the Lamb3-LKO mice (Figure 5, A and C), contrary to the reduced BEC proliferation in the Klf5-LKO mice^10^.

**Figure 5.**
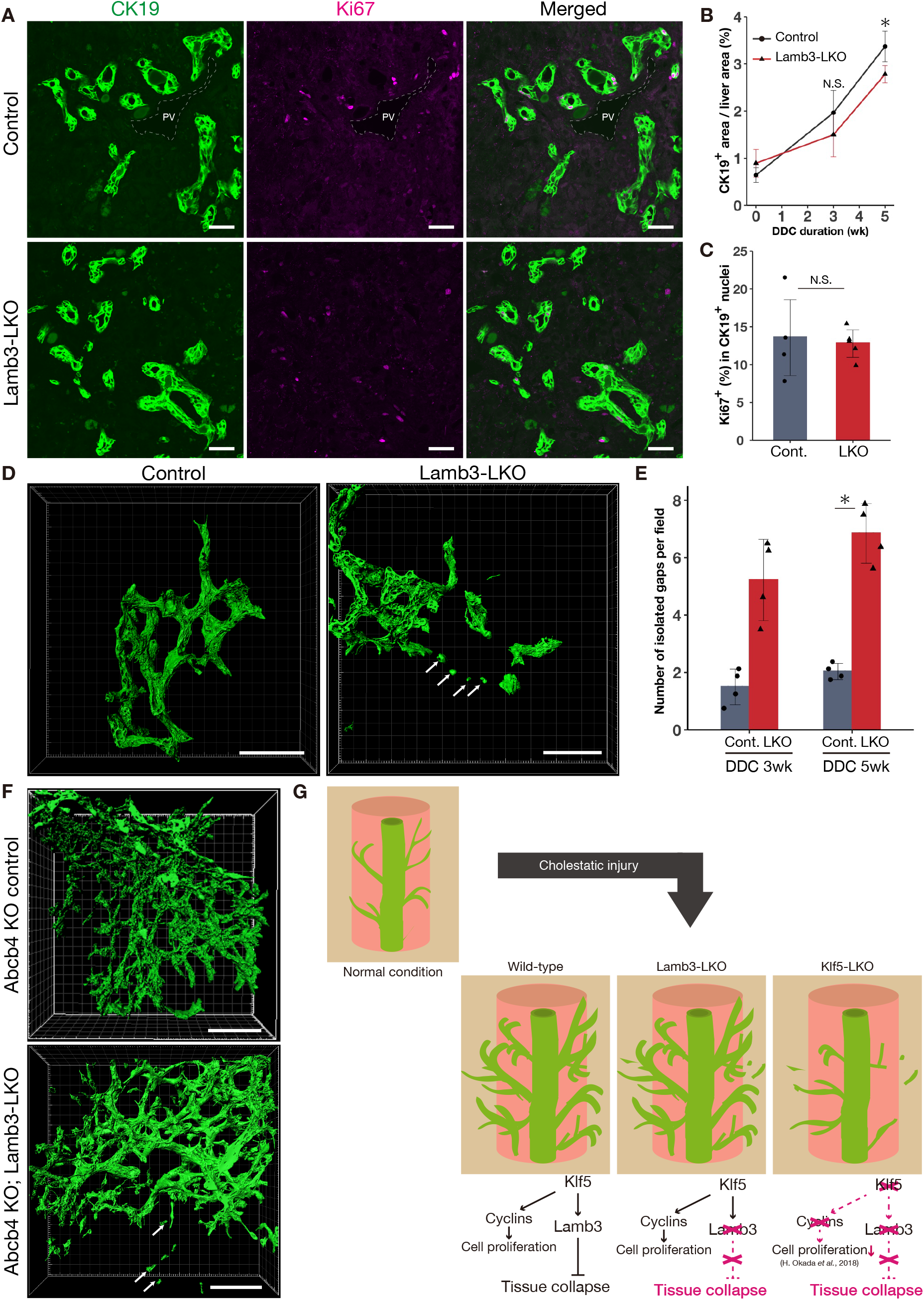
Lamb3 suppresses biliary epithelial tissue collapse under cholestatic injury conditions *in vivo*. (A) Representative images of immunostaining for CK19 (green) and Ki67 (magenta) in liver sections of control (*Lamb3 fl/fl*) and Lamb3-LKO mice treated with DDC for 3 weeks. PV, portal vein. Scale bar, 100 μm. (B) Quantification of biliary epithelial tissue expansion. Relative size of CK19^+^ areas per whole-liver section was calculated based on the immunostaining results. Data represent the mean ± S.D. For each time point, n = 5 and n = 4–6 of control and Lamb3-LKO mice were analyzed, respectively. The p-values were calculated using the two-sided Mann-Whitney U test for each time point comparing the control and Lamb3-LKO mice. An asterisk indicates that the p-value is <0.05. (C) Quantification of biliary epithelial cell proliferation. Ki67^+^ proliferating cells in CK19^+^ BEC population were counted based on the immunostaining results. Data represent the mean ± S.D. for the control (n = 4) and Lamb3-LKO (n = 5) mice. The p-value was calculated using the two-sided student’s *t*-test. (D) Representative images of the biliary epithelial tissue morphology (green) in control and Lamb3-LKO livers treated with DDC for 3 weeks as revealed by 3D-immunostaining for CK19. Stacked images were obtained with confocal microscopy and processed to reconstruct surface images using the IMARIS software. White arrows indicate CK19^+^ cell clusters separated from the biliary tree structures. Scale bars, 100 μm. (E) Quantification of the biliary epithelial tissue collapse. The numbers of isolated gaps between a detached cell cluster and the biliary branching structure were counted in livers from control and Lamb3-LKO mice after DDC treatment for 3 weeks and 5 weeks. Data represent the mean ± S.D. of n = 4 mice per genotype for each of the time points. For each mouse, 8 different fields were observed and the average number of the isolated gaps per field was calculated and presented. The p-values were calculated by the two-sided Mann-Whitney U test. An asterisk indicates that the p-value is <0.05. (F) Representative images of the biliary epithelial tissue morphology (green) in the *Abcb4* and *Lamb3* double-knockout (Abcb4 KO; Lamb3-LKO) and control (Abcb4 KO; *Lamb3 fl/fl*) livers at 8 weeks after birth. Stacked images were obtained with confocal microscopy and processed to reconstruct surface images using the IMARIS software. White arrows indicate CK19^+^ cell clusters separated from the biliary tree structures. Scale bars, 100 μm. (G) Schematic summary of the biliary epithelial tissue phenotypes observed in the Lamb3-LKO and Klf5-LKO mice in comparison to the wild-type mice under the cholestatic injury conditions. In addition, the relevant cellular functions and intracellular signaling pathways acting in BECs are illustrated.

Finally, the effect of the Lamb3 loss on the structural integrity of the intrahepatic biliary epithelia undergoing cholestasis-induced tissue remodeling was evaluated using three dimensional (3D) immunostaining analysis of the Lamb3-LKO liver. As expected, CK19^+^ cell clusters separated from the contiguous biliary tree were observed in the knockout mouse livers but not in the control; this finding strongly supported the idea that the loss of Lamb3 phenocopied the tissue collapse phenotype initially found under the Klf5-deficient condition (Figure 5, D and E). We also analyzed the liver tissues derived from a different cholestasis model based on the Abcb4 deficiency and revealed that the BEC clusters apart from bile ducts were substantially induced in the *Abcb4* and *Lamb3* double-knockout (Abcb4 KO;Lamb3-LKO) mice (Figure 5F). Taken together, these results indicate that Lamb3 is the key downstream component of Klf5 functioning independently of the pro-proliferative axis and strongly implicate Lamb3-containing laminin-332 in the maintenance of the biliary epithelial tissue integrity against cholestatic injury conditions (Figure 5G).

The ductular reaction functions as an adaptation of bile ducts via morphological alterations to ameliorate types of liver injury conditions and facilitate the organ regeneration, during which the biliary epithelia undergo multiple biological processes such as extension and sprouting of branches and increment of lumen size. In this study, visualization of the mode of biliary collapse *in vitro* and *in vivo* presented us with a hitherto unrecognized insight into the cellular function relevant to the ductular reaction. In skin epithelia, laminin-332 plays a role in adhesion to the basal lamina as a component of the hemidesmosome as well as furnishes the skin epithelial cells a movement for skin lesion compensation^28–30^. Hence, the biliary maintenance mechanism via laminin-332 seems reasonable in that it renders both flexibility and integrity to the biliary epithelia while undergoing unstable processes of tissue remodeling. Interestingly, the effect of the tissue collapse *per se* on the entire biliary tissue expansion was modest in the Lamb3-LKO mice (Figure 5G), and was much more prominent in the Klf5-LKO mice where concurrent suppression of the BEC proliferation occurred. The Lamb3-dependent biliary tissue maintenance activity may thus play an indispensable role, particularly under severely distressed cholestatic conditions, and the potential interplay between the Lamb3-containing laminin complex and growth suppressive conditions needs to be elucidated to fully understand the mechanism whereby the ductular reaction contends with devastating liver diseases.

To date, the causes of bile duct loss in human diseases, such as vanishing bile duct syndrome, have been ascribed to immunoreactions and toxic chemical components accompanying severe liver injury^31^. To the best of our knowledge, the present study is the first to demonstrate the presence and molecular basis of the tissue-autonomous biliary maintenance mechanism that counteracts the collapse of their structure upon liver injury, providing a novel therapeutic potential for hepatobiliary disorders.

## Methods

### Animals and animal experiments

All animal experiments were conducted in accordance with the Guidelines for the Care and Use of Laboratory Animals of the University of Tokyo, under the approval of the Institutional Animal Experiment Ethics Committee of the Institute for Quantitative Biosciences, The University of Tokyo (approval numbers 2804, 2904, 3004, 3004-1, 3105 and 0209). All animals were maintained under specific pathogen-free conditions. Male and female mice were used for the analyses.

Wild-type C57BL/6J mice were purchased from CLEA Japan, Inc. (Tokyo, Japan). Mice with a conditional (floxed) allele for *Klf5*^32^, Klf5-LKO mice^10^, and Lamb3-LKO mice^24^ were reported previously and maintained on a C57BL/6 background. Abcb4 KO mice^26^ (FVB.129P2-Abcb4<tm1Bor>/J, Stock No. 002539) were purchased from Jackson Laboratory (Bar Harbor, ME, USA) and backcrossed to C57BL/6J mice for at least 10 generations. For the DDC injury model, mice were fed a diet containing 0.1% DDC (F-4643; Bio-Serv) for the period indicated.

For the hepatocyte-specific labeling and tracing experiments, the recombinant adeno-associated virus expressing a histone H2B-mVenus fusion reporter under the control of the ubiquitous CAG promoter (rAAV2/8-H2B-mVenus) was packaged in HEK293 cells according to a protocol described previously^10,15^. The titered virus was delivered to adult Klf5-LKO mice by intraperitoneal injection at a dose of 1×10^11^ vector genomes/mouse. Two weeks after infection, the mice were fed the DDC diet for two and a half weeks.

### Immunostaining analysis of liver tissue sections and quantification

Dissected mouse livers were directly embedded in Tissue-Tek O.C.T. Compound (4583; Sakura Finetek USA, Inc.) and snap-frozen. Frozen sections (8 µm) of the liver were prepared using an HM525 cryostat (Microm International) and placed on aminopropyltriethoxysilane-coated glass slides (Matsunami Glass). Fixation was performed with either acetone and/or 4% paraformaldehyde after sectioning, according to the optimized conditions for individual primary antibodies, as described in Supplementary Table 1. After blocking in 3% fetal bovine serum (FBS) in PBS containing 0.1% Triton X-100, the samples were incubated with primary antibodies at dilutions as described in Supplementary Table 1 and then with fluorescence-conjugated secondary antibodies. Nuclei were counterstained with Hoechst 33342 (Sigma-Aldrich). Images were obtained with a confocal laser-scanning microscope (Fluoview FV3000, Olympus), except for the images in Supplementary Figure S4 that was obtained using an epifluorescent microscope (Axio Observer.Z1, Zeiss). For 3D presentation, surfaces were virtually constructed using the “Surface” function in IMARIS software (Bitplane).

Quantitative evaluations of CK19^+^ area in the liver sections and Ki67^+^ cells in CK19^+^ cells were done as previously described^10^. For manual counting of the isolated spot numbers in Figure 5E, 3D images were acquired with a spinning-disk confocal microscope (SpinSR10, Olympus).

### Sorting and culture of mouse BECs

Preparation of EpCAM^+^ cells from adult mice was performed as previously described^19^. In brief, a cell suspension from adult mouse livers was prepared using a two-step collagenase perfusion method, and the parenchymal cell (i.e., hepatocyte) and non-parenchymal cell (NPC) fractions were prepared by centrifugal separation. To prepare the BEC fraction, NPCs were treated with anti-EpCAM monoclonal antibody, and the samples were sorted using Moflo XDP (Beckman Coulter). Non-viable cells were excluded through propidium iodide staining. EpCAM^+^ cells were seeded on a type-I collagen-coated dish and incubated in the culture medium as previously reported^19^ (Williams’ medium E containing 10% FBS (JRH), 10 mM nicotinamide (Sigma), 2 mM L-glutamine (Gibco 25030-081), 0.2 mM ascorbic acid (Sigma, A8960), 20 mM HEPES pH7.5 (Gibco), 1 mM sodium pyruvate (Sigma, S8636), 17.6 mM NaHCO_3_ (Wako, 191-01305), 14 mM glucose (Wako, 041-00595), 100 mM dexamethasone (Sigma), 1× insulin/transferrin/selenium (Gibco), 10 ng/mL hepatocyte growth factor, 10 ng/mL mouse epidermal growth factor, and 50 mg/mL gentamicin). EpCAM^+^ cells sorted from the *Klf5*^*flox/flox*^ mice without the *Alfp-Cre* transgene were cultured for approximately one month until the cells constantly expanded, and cell lines were established after a few passages and used for experiments.

### Establishment of the modified BEC lines

To establish the inducible *Klf5* knockout BEC line, a pMXs-based retroviral vector was used to introduce and express the CreERT2-IRES-mVenus cassette under the control of the MMLV LTR promoter. Plat-E cells^33^ seeded on a 10cm-culture dish were transfected with the vector plasmid using PEI Max (Polysciences, 24765) according to the manufacturer’s recommendations, and the culture medium was changed to the one used for culturing BECs described above, 24 h after transfection. Forty-eight hours after transfection, the culture medium was collected and centrifuged, and the supernatant was purified through filtration. The BEC line harboring *Klf5*^*flox/flox*^ was cultured for two days with the purified medium for retroviral infection, and cells stably expressing the introduced gene cassette were sorted using Moflo XDP (Beckman-Coulter) based on mVenus fluorescence as an indicator. The sorted cells were cultured and expanded, referred to as the inducible *Klf5* knockout (Klf5-iKO) cell line. For inducible knockout experiments, the culture medium was supplemented with 4-OHT at a final concentration of 0.1 µM.

To establish an inducible Klf5-overexpressing BEC line, a piggyBac transposon system was used as previously described^34^. A piggyBac transposon vector containing Klf5 under the Tet Response Element (TRE) was constructed from the pB-TRE-dCas9-VPR^35^ and 4xmts-mScarlet-I^36^ vectors (purchased from Addgene as plasmids #63800 and #98818, respectively). The piggyBac transposon vector and pBase transposase-containing vector were transfected into Klf5-iKO cells using PEI Max. The cells stably expressing the introduced gene cassette were sorted using Moflo XDP (Beckman-Coulter) based on mScarlet1 fluorescence as an indicator. To achieve overexpression of Klf5, the culture medium was supplemented with doxycycline (Dox) to attain a final concentration of 1 µg/mL.

### The 3D culture of BECs for biliary branching structure formation

To achieve the formation of the biliary tissue-like branching structure *in vitro*, BEC lines were cultured using a modified protocol based on a method previously described^19^. In brief, the type I collagen mix solution was prepared by mixing a 20:4:3:3 ratio of 0.3% Cellmatrix Type I-A (Nitta Gelatin), 10× DMEM/ F12 (Sigma), 1× PBS, and the reconstitution buffer (50 mM NaOH, 200 mM HEPES pH7.5, 262 mM NaHCO_3_). The bottom layer of the culture was first prepared by pouring 100 μL of type I collagen mix solution into each well of an 8-well coverglass chamber (177402PK, Nunc, Roskilde, Denmark). The middle layer of the culture was prepared by adding 100 μL of the type I collagen mix solution containing 3 × 10^3^ cells/well of the Klf5-iKO line and laid over the bottom layer. The culture medium comprising DMEM/F-12, 10% FBS (Gibco), 100 mM dexamethasone (Sigma), 1× insulin/transferrin/selenium (Invitrogen), 50 mg/mL gentamicin, 10 ng/mL hepatocyte growth factor, and 10 ng/mL mouse epidermal growth factor was gently laid over the gel layers. Biliary branching-like structures can grow for at least 10 d. In all the experiments in this study, 4-OHT was added seven days after embedding Klf5-iKO cells. Notably, the number of embedded cells and culturing terms are important for mimicking the biliary branching-like structure. The culture medium was changed every three days.

### Gene expression analysis by quantitative RT-PCR

*In vitro*-formed biliary branching structures were collected from the collagen-embedded cultures by collagenase treatment. Total RNA was isolated using the TRIzol reagent (Invitrogen), treated with DNase I (Invitrogen), and then used for cDNA synthesis with PrimeScript RT Master Mix (Takara). Quantitative PCR analyses were performed using the LightCycler 96 System (Roche Applied Science) with SYBR Premix Ex Taq (Takara). The mouse *Rplp0* gene was used as an internal control. The primer sequences are listed in Supplementary Table2.

### Image acquisition and quantification of *in vitro* branching structures

The *in vitro*-formed biliary branching structures were observed using a fluorescence phase-contrast microscope (BZ-X800, Keyence). The 8-well coverglass chambers were kept incubated at 37°C with 5% CO2. Images of the individual regions of interest were taken at the indicated time points in each panel. For quantification, each view was taken every 30 min from 24 to 48 h after EtOH (control) or 4-OHT administration. Three to seven views of each well were randomly chosen, and four to five individual wells of each group were quantified. The number of nascent spots was manually counted.

### EdU assay

To monitor the level of BEC proliferation in the *in vitro* culture, EdU-labeling assays were performed using the Click-iT Plus EdU AlexaFluor488 cytometry assay kit (Life Technologies, Inc.), according to the manufacturer’s instructions. Forty-eight hours after 4-OHT administration, EdU was added to the culture media at a final concentration of 10 µM for 2 h. The samples were analyzed using Moflo XDP (Beckman Coulter) with non-viable cells excluded using the Fixable Viability Stain 450 (BD Biosciences).

### RNA-seq

*In vitro* biliary branching structures were collected from the collagen-embedded cultures by collagenase treatment at 24 and 48 h after EtOH (control) or 4-OHT administration. Total RNA was isolated using the TRIzol reagent (Invitrogen), treated with DNase I (Invitrogen), and purified using the TRIzol reagent. The quality of the collected RNAs was examined using a Bioanalyzer (Agilent), and their RINs were confirmed to be over 8.0. cDNA libraries were prepared from 500 to 1000 ng of total RNA using the TruSeq Stranded mRNA Library Prep kit (Illumina) according to the manufacturer’s instructions and sequenced on a HiSeq 2500 (Illumina).

### Bioinformatic analysis of RNA-seq and ChIP-seq data

The first 14 bases from each read were trimmed, and the subsequent 52 bases were aligned to the *Mus musculus* genome (Gencode v22) using Hisat2. The generated SAM files were converted to BAM files using Samtools v1.8^37^. Only the reads that were uniquely aligned to the transcripts were counted. Transcript counts were normalized, and differential gene expression was calculated using the DESeq2^38^ package in R software version v.3.6.1, assisted by the Integrated Development Environment for R v.1.2.1335 (https://www.rstudio.com/). Significant DEGs were selected based on a false discovery q-value cutoff of 0.05. Gene enrichment analysis of DEGs was performed using Metascape^39^.

To examine the direct transcriptional target genes of mouse Klf5 and human KLF5, average MACS2 binding scores on the upstream sequence within 1 kbp from the transcriptional start site of potential Klf5 targets identified through RNA-seq (the intersection of the three circles in Figure 3B) were accessed by publicly available ChIP-seq datasets via ChIP-atlas (http://chip-atlas.org)^21^ using *Mus musculus* (mm10) and *Homo sapiens* (hg19) genomes, respectively.

### Statistics and reproducibility

In all animal and cell culture experiments, the samples represent biological replicates derived from different mouse and culture well individuals, respectively. Data are expressed as the mean ± S.D. The Shapiro-Wilk test was used to assess the normality of the distribution of the investigated parameters, and significant differences were tested using the unpaired two-sided Mann-Whitney U test or Student’s t-test accordingly. Statistical analyses were performed using the R software and the Prism software (GraphPad, San Diego, CA, USA). Differences were considered statistically significant at p < 0.05.

## Data availability

The RNA-seq data (FASTQ files) were deposited in the NCBI database under the accession number GSE145420. Analysis of this dataset used standard bioinformatics tools and codes (as described in the section titled **Bioinformatic analysis of RNA-seq and ChIP-seq data**), which are available upon request.

## Acknowledgments

We thank Prof. K. Kaestner for providing the Alfp-Cre transgenic mice; Prof. R. Nagai (Jichi Medical University, Tochigi, Japan) and Prof. I. Manabe (Chiba University, Chiba, Japan) for the anti-Klf5 antibody; Prof. I. Alexander (Children’s Medical Research Institute, Australia) for the pAM-LSP1-EGFP plasmid; Prof. R. J. Samulski and the NGVB Biorepository (University of North Carolina at Chapel Hill, NC) for the XX6-80 plasmid; Prof. R. Zeller for the pDIRE plasmid; and Penn Vector Core (University of Pennsylvania) for the p5E18-VD2/8 plasmid. We also thank Drs. K. Fujiki and K. Nakagawa and Prof. K Shirahige (Institute of Molecular and Cellular Biosciences, University of Tokyo, Japan) and M Muraoka (National Institute of Genetics, Japan) for help with RNA-sequencing; A Maeno for help with video editing (National Institute of Genetics, Japan); C. Koga for help with cell sorting; and The University of Tokyo IQB Olympus Bio-imaging Center (TOBIC) for help with microscopy and image acquisitions. The TROMA-III, developed by Dr. Rolf Kemler, was obtained from the Developmental Studies Hybridoma Bank, developed under the auspices of the NICHD and maintained by the Department of Biology at The University of Iowa.

This work was supported by PRIME from the Japan Agency for Medical Research and Development (AMED) JP21gm6210001 (to T.I.) and by the Japan Society for the Promotion of Science KAKENHI Grants 17J06605 (to YM) and 21H02518 (to TI).

## Author contributions

HO, conception and design, acquisition of data, analysis and interpretation of data, drafting the article, contributed unpublished essential data or reagents; MY, design and acquisition of data, analysis and interpretation of data, contributed unpublished essential data or reagents; ME, contributed essential reagents; YK, contributed essential reagents; AM, drafting the article; TI, conception and design, analysis and interpretation of data, drafting the article. All authors reviewed the results and approved the final version of the manuscript.

## Supplementary Information

**Supplementary Figure 1. Focal plane analysis by image processing to evaluate the absence of co-localization of the isolated CK19+ cells and the hepatotropic H2B-mVenus expression**.

(A) Representative 3D images of the rAAV2/8-H2B-mVenus-infected liver sections after DDC administration as observed by confocal microscopy. Biliary epithelial tissue structure (CK19 immunostaining, green) and H2B-mVenus signals (magenta) are shown. The white arrows indicate the isolated CK19^+^ cells and clusters from bile ducts observed in the Klf5-LKO livers. Scale bars, 100 μm. (B) The raw original image shown in A for the Klf5-LKO liver (left panel; lateral view of the same 3D image is also included) was first processed to acquire surfaces of the biliary epithelial tissue structure and H2B-mVenus^+^ nucleus (middle panel), and then processed so that the H2B-mVenus signals out of the Z-plane where the isolated CK19+ cells reside were eliminated (right panel).

**Supplementary Figure 2. Establishment and characterization of Klf5-iKO cell line**.

(A) Workflow of the establishment of a mouse biliary epithelial cell line with tamoxifen-inducible *Klf5* gene knockout (*Klf5 flox/flox*;*LTR-CreERT2-IRES-mVenus*, Klf5-iKO).

(B) Time course of the reduction in *Klf5* expression upon tamoxifen administration in the *in vitro* culture system of the Klf5-iKO cells. The cells were cultured in the presence of tamoxifen (4-OHT) or EtOH (control) for the indicated periods and the gene expression levels were quantitatively analyzed. Data represent the mean ± S.D. of *n* = 4 biological replicates per time point for each treatment.

(C) Representative bright-field images of the *in vitro* culture system of the wild-type biliary epithelial cell line. The cells were cultured for 7 days to form branched biliary organoids and then for an additional 2 days in the presence of tamoxifen (4-OHT) or EtOH (shown as control), showing that the tamoxifen treatment per se did not result in the collapse of the biliary organoid structures. Scale bars, 100 μm.

**Supplementary Figure 3. Cellular and genetic alterations induced under the collapse of the expanding biliary epithelial tissue *in vitro***.

(A and B) MA plots showing differentially expressed genes between 4-OHT-treated and control conditions in Klf5-iKO cells in the *in vitro* culture system at 12 (A) and 24 (B) hours after 4-OHT administration. The common responsible genes that were identified and shown in bold in Figure 3D are highlighted in yellow color. The other DEGs and non-DEGs are shown in black and grey, respectively.

(C and D) Effect of tamoxifen-induced Klf5 knockout on the proliferation of Klf5-iKO cells in the *in vitro* culture system. The level of cell proliferation was analyzed using the EdU incorporation assay and flow cytometric analysis at 48 h after 4-OHT administration. (C) Representative scatter plot images of flow cytometric analysis. (D) Quantitative evaluation of the ratio of EdU+ cells. Data represent the mean ± S.D. of *n* = 4 biological replicates for each condition. Statistical significance was evaluated using the two-sided Student’s *t*-test to be not significant (N.S.).

(E) The mean MACS2 score (http://chip-atlas.org) calculated for potential targets of human KLF5 in human cases.

(F and G) Klf5/KLF5 binding sites around the *Lamb3*/*LAMB3* transcription start sites in mouse (F) and human (G) genomes revealed by Peak Browser of ChIP-atlas. Displayed transcripts correspond to ENSMUST00000016315.15 (mouse) and ENST00000356082.8 (human). Vertically long rectangles indicate coding sequences (CDS). Blue, green and red color bars indicate a relative binding probability in an ascending order.

**Supplementary Figure 4. Cholestasis-induced deposition of Lama3 around the expanding biliary epithelia was suppressed in Klf5-LKO mouse livers**.

(A) Representative images of immunostaining for Lama3 (shown in magenta and white colors in the upper and lower panels, respectively) and CK19 (green in the upper panels) in liver sections of Klf5-LKO and control (*Klf5 fl/fl*) mice treated with DDC for 3 weeks. Loss of Lama3 expression in the Klf5-LKO mouse liver was confirmed at the protein level. Scale bars, 100 μm.

(B) Representative images of immunostaining for Lama3 (shown in magenta and white colors in the upper and lower panels, respectively) and CK19 (green in the upper panels) in liver sections of *Abcb4* and *Klf5* double knockout (Abcb4 KO;Klf5-LKO) and control (Abcb4 KO;*Lamb3 fl/fl*) mice at 8 weeks after birth. Scale bars, 100 μm.

**Supplementary Figure 5. Cholestasis-induced deposition of the laminin-332 components around the expanding biliary epithelia was diminished in Lamb3-LKO mouse livers**.

(A) Representative images of immunostaining for Klf5 (magenta and white in the upper and lower panels, respectively) and CK19 (green) in liver sections of Lamb3-LKO and control (*Lamb3 fl/fl*) mice under the physiological conditions in the absence of liver injury. Scale bars, 100 μm.

(B–E) Representative images of immunostaining for Lamb3 (B), Klf5 (C), Lama3 (D), and Lamc2 (E) in liver sections of Lamb3-LKO and control (*Lamb3 fl/fl*) mice treated with DDC for 3 weeks. The signals for the proteins of interest are shown in merged images (upper panels, magenta) with CK19 co-staining (green), as well as in single-channel images (lower panels, white) for ease of visibility. Scale bars, 100 μm.

**Supplementary Table 1.**
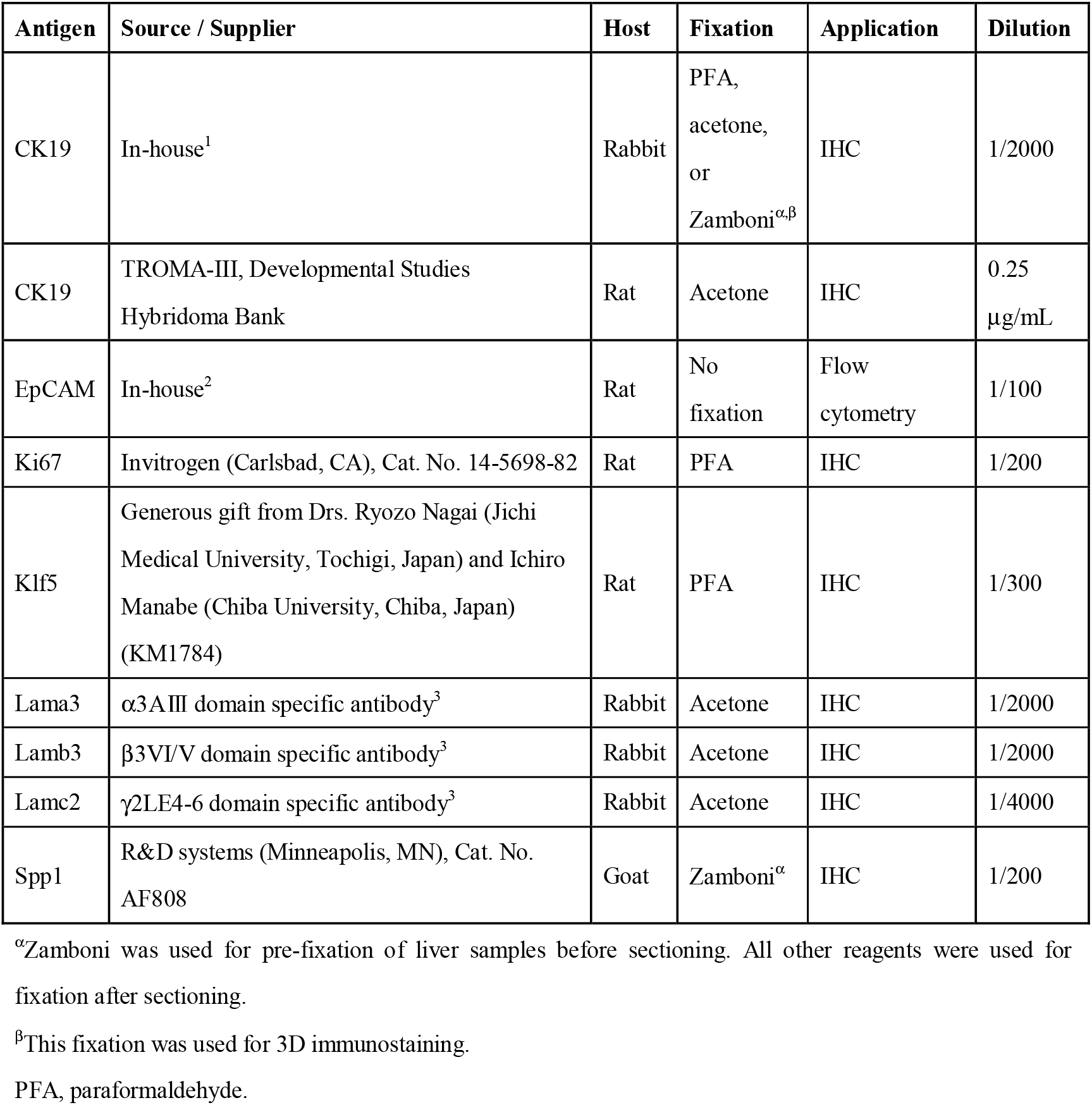
Antibodies used in this study.

**Supplementary Table 2.**
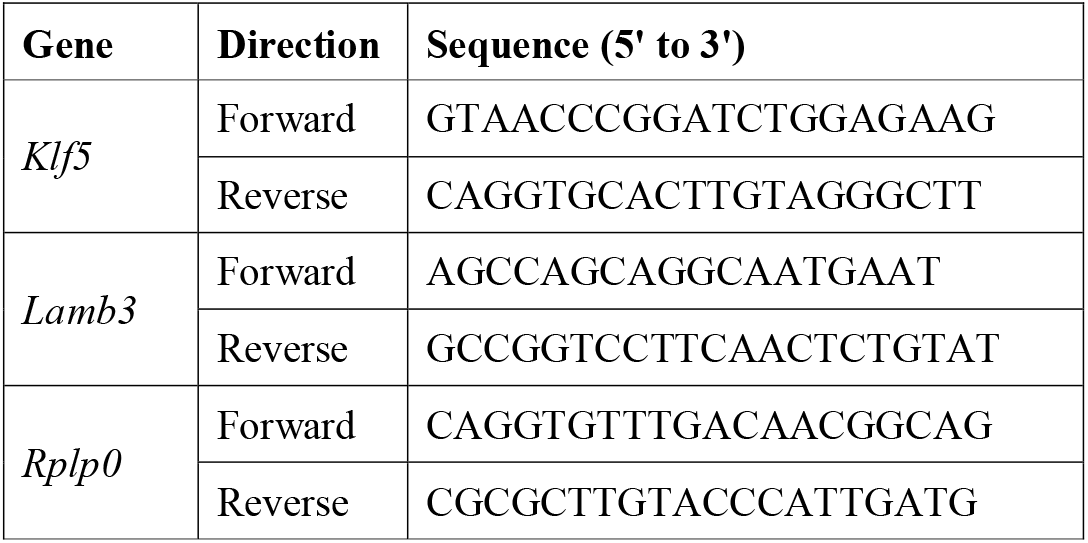
List of primers used in qPCR for detecting mRNAs in this study.

