## Supplementary figures and images for "KLF5-regulated extracellular matrix remodeling secures biliary epithelial tissue integrity against cholestatic liver injury"

### Figure_bioRxiv_sup.pdf

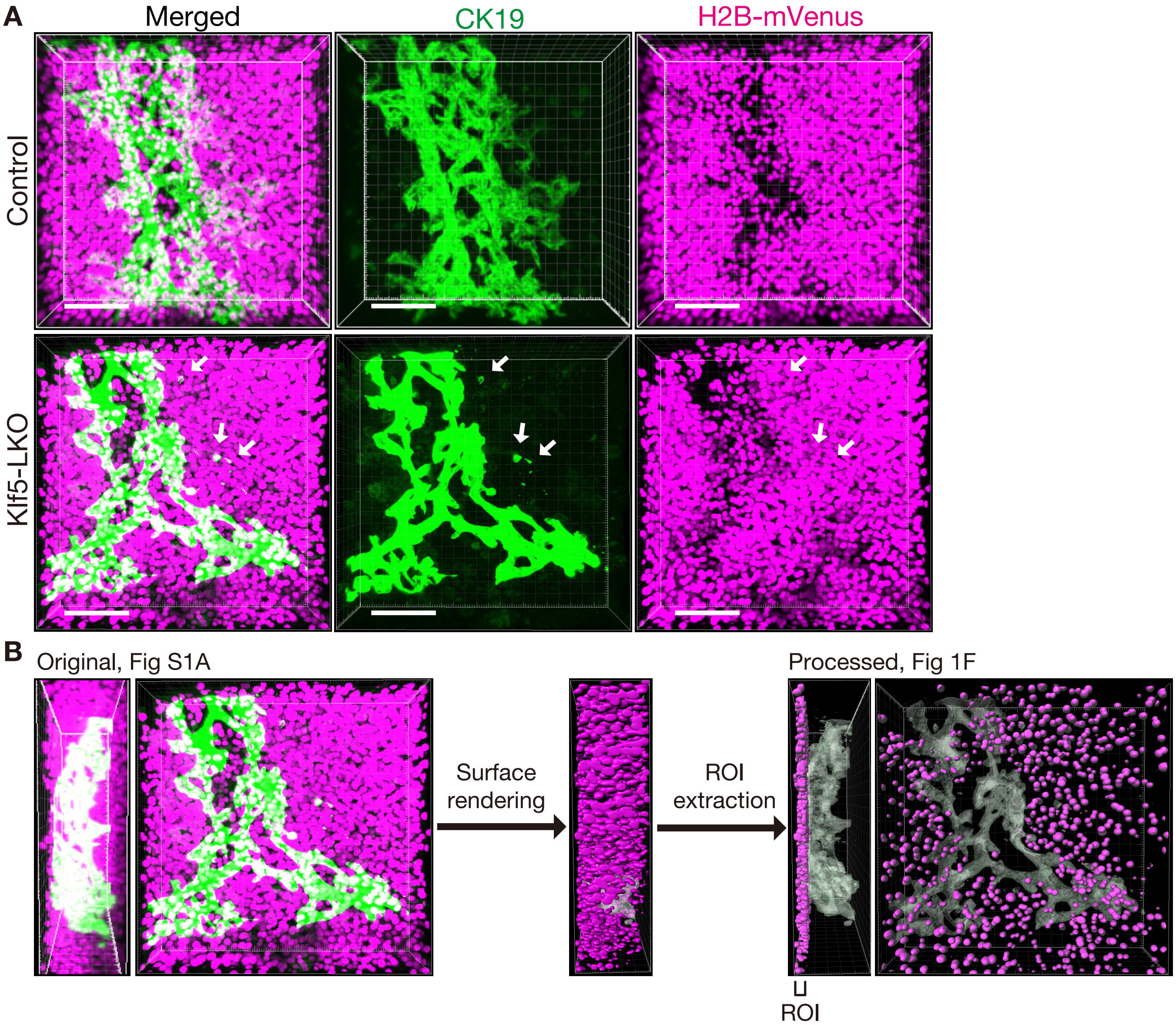

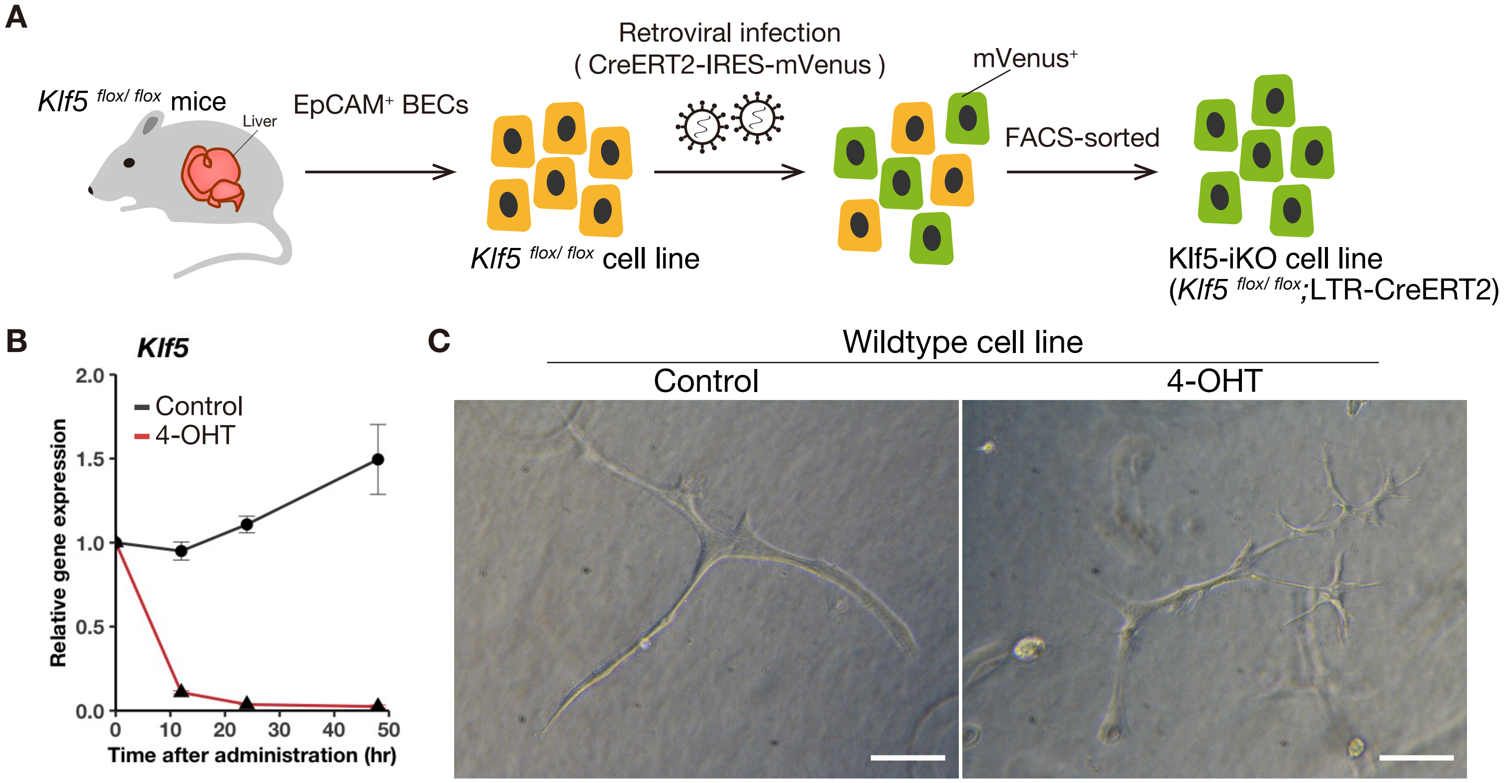

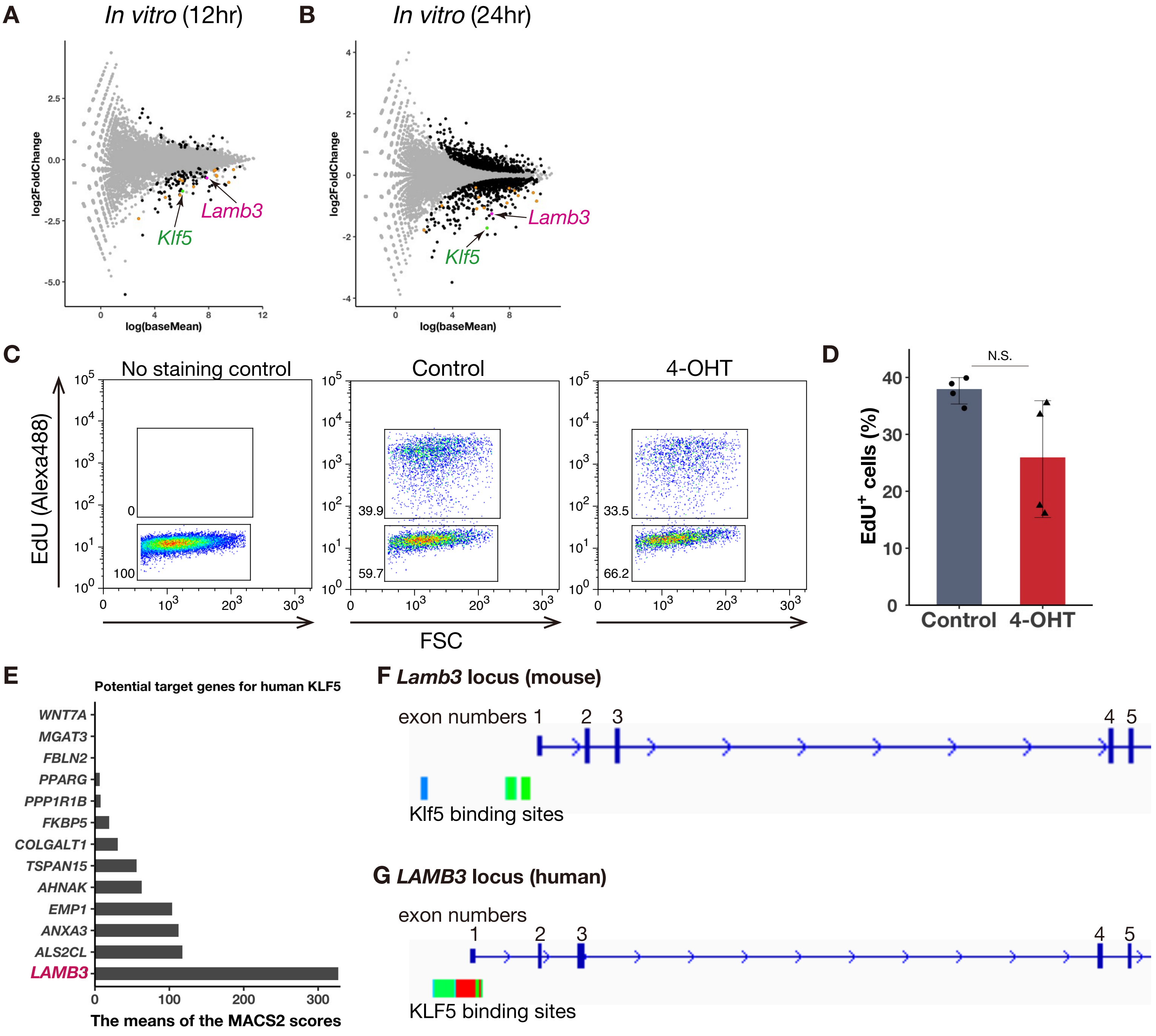

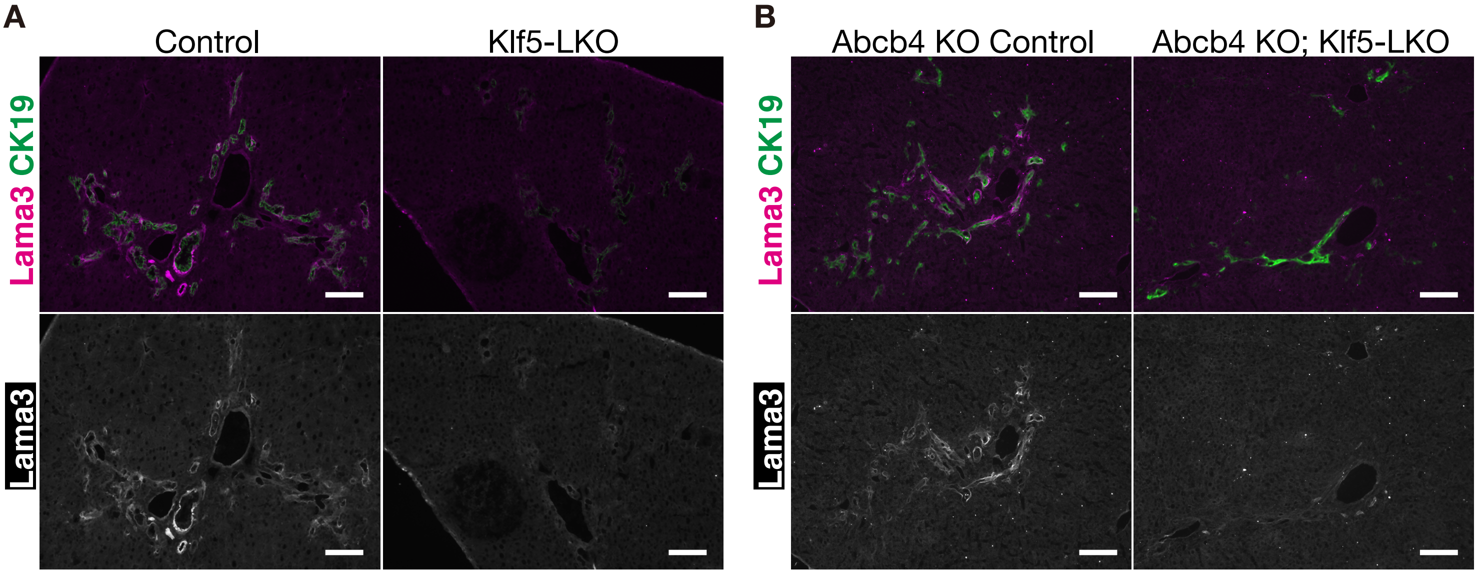

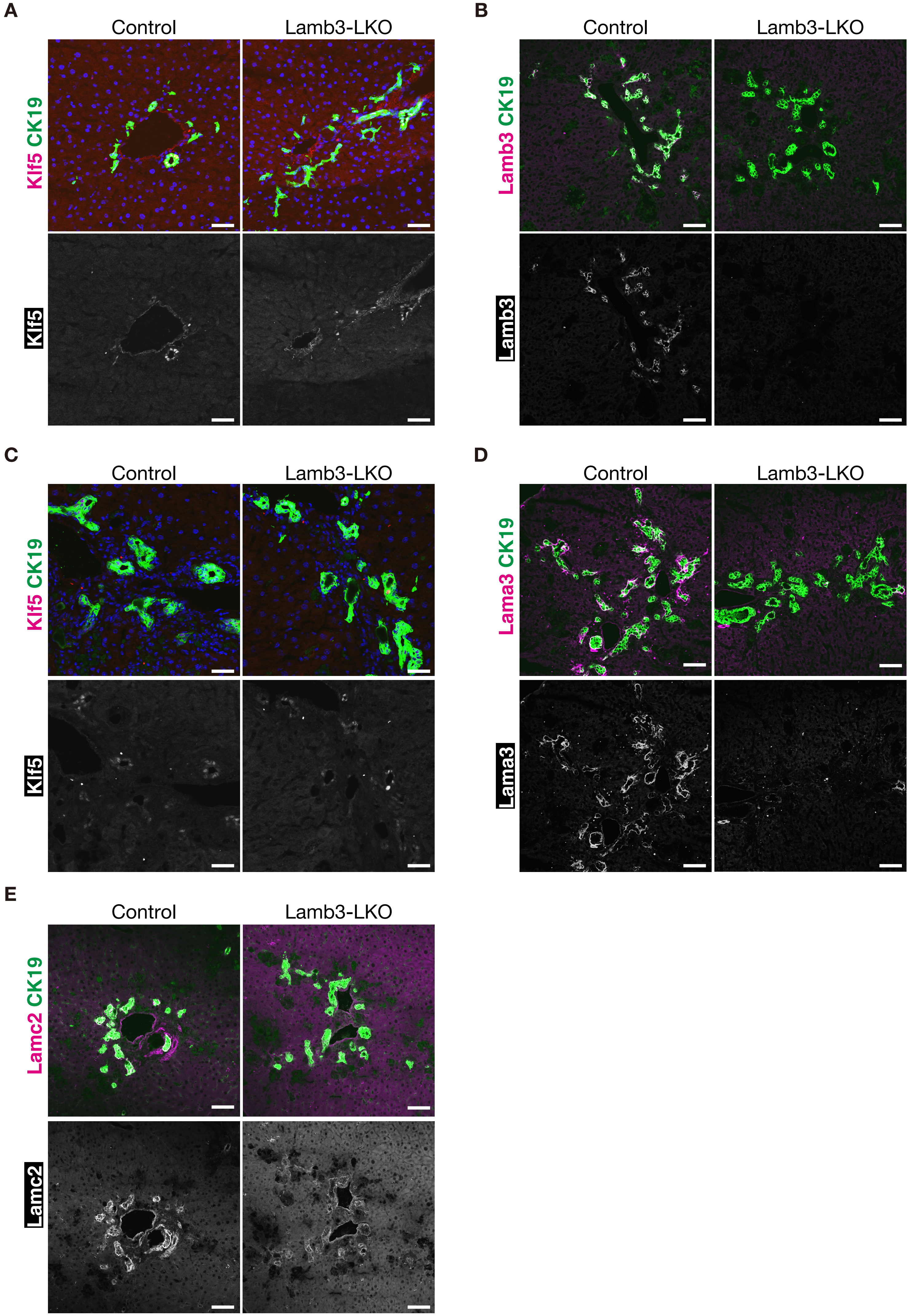
